# Cortical Processing for the Vestibular and Visual Input of Egomotion in Macaque Monkeys: Separate Networks with Targeted Convergence

**DOI:** 10.1101/2025.08.06.668905

**Authors:** Sarah Marchand, Vanessa De Castro, Elisabeth Excoffier, Marie-Alphée Laurent, Maxime Rosito, Nathalie Vayssière, Benoit R. Cottereau, Alexandra Séverac Cauquil, Jean-Baptiste Durand

## Abstract

The integration of visual and vestibular inputs during egomotion is fundamental for both postural and navigational control. In the present study, we used functional magnetic resonance imaging (fMRI) in four macaque monkeys to investigate cortical activation in response to galvanic vestibular stimulation (GVS), applied through transmastoid electrodes, and egomotion-compatible (EC) optic flow patterns. Visual and vestibular stimulations activate two largely independent cortical networks: the vestibular network encompasses the insular cortex, superior parietal lobule, frontal lobe, and cingulate cortex, while optic flow primarily activates regions in the superior temporal sulcus, temporo-parietal junction, inferior parietal lobule, and restricted portions of the cingulate and frontal cortices. Despite this segregation, several areas exhibit visuo-vestibular convergence: VPS in the temporo-parietal junction, area 7 in the inferior parietal lobule, VIP and LIP in the intraparietal sulcus, MSTd in the superior temporal sulcus, CSv in the cingulate sulcus, and FEFsem in the frontal cortex. These findings demonstrate that visual and vestibular signals generated by egomotion are processed in extended and distinct cortical networks with several narrow convergence sites, consistent with the idea that multisensory integration during self-motion is achieved through selective convergence rather than general network overlap.

## 1. Introduction

Visuo-vestibular integration during egomotion represents a fundamental process for postural and navigational control in human and non-human primates. While both sensory modalities are activated during egomotion, we still have a fragmented view about how and where they converge at the cortical level. The vestibular system provides sensory foundation for detecting egomotion by its peripheral components: three orthogonally oriented semicircular canals to detect rotation of the head in space, and two otolith organs (utricle and saccule) to detect linear accelerations (Iurato, 1967; Kingma & van de Berg, 2016). Together, these systems form a robust tool for detecting gravitational forces and both angular and linear head motion necessary for spatial orientation and balance (Goldberg *et al*., 1984; Day & Fitzpatrick, 2005). Vestibular input provides immediate, unambiguous signal of head rotation and acceleration, with reaction times of the order of milliseconds following movement onset (Fernandez & Goldberg, 1971; Eatock *et al*., 2008). When vestibular function is impaired, individuals suffer spatial disorientations and navigation difficulties (Brandt & Dieterich, 1999; Seemungal *et al*., 2007), which provides indirect evidence for the essential role of the vestibular system in self-motion perception.

To study vestibular processing, galvanic vestibular stimulation (GVS) is now the method of choice due to its ability to stimulate the whole vestibular system (Marchand, Langlade, Legois *et al*., 2025 for a recent review). GVS consists of sending a low-intensity electric current through electrodes stuck on the skin at the mastoid process level to directly activate vestibular neuron afferents. On the anodal side, there is a decrease in neuronal activity, whereas cathode increases it, and this discrepancy generates ocular torsion (Séverac Cauquil *et al*., 2003) and body tilt (Nashner & Wolfson, 1974), directed toward the anode, and an illusion of egomotion can be reported (Fitzpatrick & Day, 2004). This non-invasive method opens a way to study vestibular processes and is particularly useful in neuroimaging research since it can be combined with functional magnetic resonance imaging (fMRI). Despite the fact that vestibular processing undoubtedly plays an essential role in the integration of egomotion, gaps in knowledge regarding cortical vestibular networks remain. Human neuroimaging studies consistently outlined a distributed vestibular network that included the dorsal portion of the medial superior temporal area (MSTd), ventral intraparietal area (VIP), cingulate sulcus visual area (CSv), and the parieto-insular vestibular cortex (PIVC) (Smith *et al*., 2018; Marchand, Langlade, Legois *et al*., 2025). However, little is known about the organization of the cortical vestibular network in non-human primates (NHPs). Important insights into macaque visuo-vestibular integration have been gained from seminal work that examined regions like the ventral posterior suprasylvian (VPS) area, the medial superior temporal (MST) area, the frontal eye field (FEF) area, and the visual area V6. Nonetheless, these studies used focal electrophysiological approaches instead of whole-brain mapping techniques (Angelaki & Cullen, 2008; Chen *et al*., 2011b; Gu *et al*., 2008, 2010, 2012). Similarly, although some recent research has investigated the neural mechanisms of GVS in NHPs, explaining how GVS activates canal and otolith afferents (Forbes *et al*., 2023; Kwan *et al*., 2019), whole-brain cortical activations have not yet been investigated. Additionally, no research has combined GVS and fMRI in NHPs to date, making it impossible to compare the two species directly and to identify the large-scale cortical networks involved in vestibular stimulation.

The visual system’s key role in egomotion perception has been studied in greater depth. Gibson’s initial study defined optic flow and presented the process by which patterns of motion within the visual field create information about self-motion through the ‘focus of expansion’ (Gibson, 1950). Following studies, including Lee’s « swinging room » experiments (Lee & Aronson, 1974; Lee & Lishman, 1975), demonstrated that postural response and perception of self-motion can be evoked by optic flow alone. Willing to examine optic flow in a strict paradigm, Smith and collaborators (Smith *et al*., 2006) developed the « 1P/9P » paradigm where egomotion-compatible (EC) single-patch stimuli were compared with egomotion-incompatible (EI) nine-patch arrays. This technique has unveiled large-scale cortical networks for visual self-motion processing including MSTd, VIP, and other areas in humans but also in non-human primates (Cottereau *et al*., 2017).

The main resulting question is how visual and vestibular networks interact with each other. While visuo-vestibular integration has been recognized in human neuroimaging (Aedo-Jury *et al*., 2020; Marchand *et al*., 2025), the precise sites and mechanisms by which it occurs in non-human primates have been fruitfully studied with methods limited to focal explorations (Angelaki & Cullen, 2008; Chen *et al*., 2011b; Gu *et al*., 2008, 2010, 2012). Deepening our understanding of this process is, however, important for basic neuroscience as well as to ascertain the translational validity of NHPs model to understand human’s postural and navigational control. The present study bridges these gaps in fundamental knowledge by combining GVS and fMRI for the first time in rhesus macaques, and by offering a comparison between vestibular activation patterns and visual egomotion-responsive areas. Our ultimate objective is to identify the cortical network activated by GVS and measure its overlap with visual egomotion-responsive areas to identify the sites of visuo-vestibular convergence. We hypothesize that visuo-vestibular integration is attained through selective convergence in some cortical regions rather than global networks overlap.

## 2. Methods

### 2.1. Ethics statement

The authors declare no conflict of interest related to this work. All procedures including non-human primates were conducted in accordance with current European legislation (Directive 2010/63/EU) and French regulations (Décret 2013–118) governing the use of animals in scientific research. The project received approval from the local ethics committee (CNREEA code: C2EA–14) and was authorized by the French Ministry of Research (authorization number: MP/03/34/10/09).

### 2.2. Subjects

Four female rhesus macaques (*Macaca mulatta*) (M01, M02, M03, and M04) (5–15 years, 5–8 kg) were involved in this study. They were housed together in a large, enriched enclosure and could thus develop natural social and foraging behaviors. They returned to their individual cages to be fed twice a day, with standard primate biscuits supplemented with various types of fruits and vegetables. Health inspections were carried out quarterly. Details about the animals’ surgical preparation are provided in Cottereau *et al*., 2017. These individuals were trained to stare at a screen in a room simulating the scanning environment, to accustom them to the task and the noise generated by the setup. They were also accustomed to the visual stimuli used in this study, to avoid any novelty effect during acquisition.

### 2.3. Experimental design

#### 2.3.1. Galvanic Vestibular Stimulation (GVS)

We used GVS to stimulate the vestibular apparatus using a low-intensity current transmitted via electrodes placed on the skin surface (Marchand, Langlade, Legois *et al*., 2025). The current generator (Digitimer, UK; model DS5) was piloted by a computer located in the MRI control room via a USB data acquisition card (NI USB-6009, DAQ 8AD 2DA 14bit 48kS/s Labview). The use of a current-limited stimulator ensured safety and stability of the stimulation, preventing any excessive current that could cause any sort of damage. The electric signal went through the filter plate of the MRI, which acts as a low-pass filter to prevent high-frequency noise from entering the MRI room and interfering with image acquisition, sending the resulting current into the electrodes. The two gold electrodes (1 cm diameter) were attached to the skin of the monkey over each mastoid, and were covered with conductive jelly to increase impedance and improve signal delivery (**Figure 1A**). Before positioning the electrodes, the subject’s skin was prepared with alcohol. The electric stimulation was delivered during periods lasting 18 seconds. The waveform was sinusoidal, to avoid sharp ON/OFF transitions, with a frequency of 1 Hz and an amplitude of ± 1.25 mA (**Figure 1B**). This frequency was chosen since 1 Hz falls within the range of vestibular afferent modulation of physiological head motion stimuli (0.1–25 Hz) for physiological head motion stimuli (Goldberg *et al*., 2012; Kwan *et al*., 2019), and sinusoidal waveforms at this frequency are known to cause strong vestibular responses in human subjects with minimal discomfort. The ± 1.25 mA amplitude was selected based on earlier research demonstrating that the general range of individual optimal current intensity is about 1-2.5 mA, seen across subjects (Caudron *et al*., 2018; for a recent review of GVS parameters, see Valter *et al*., 2025). In a similar range, Kwan *et al*. (2019) used a 2-mA peak-to-peak amplitude on awake macaque monkeys, obtaining strong and robust primary vestibular afferents and ocular responses.

**Figure 1.**
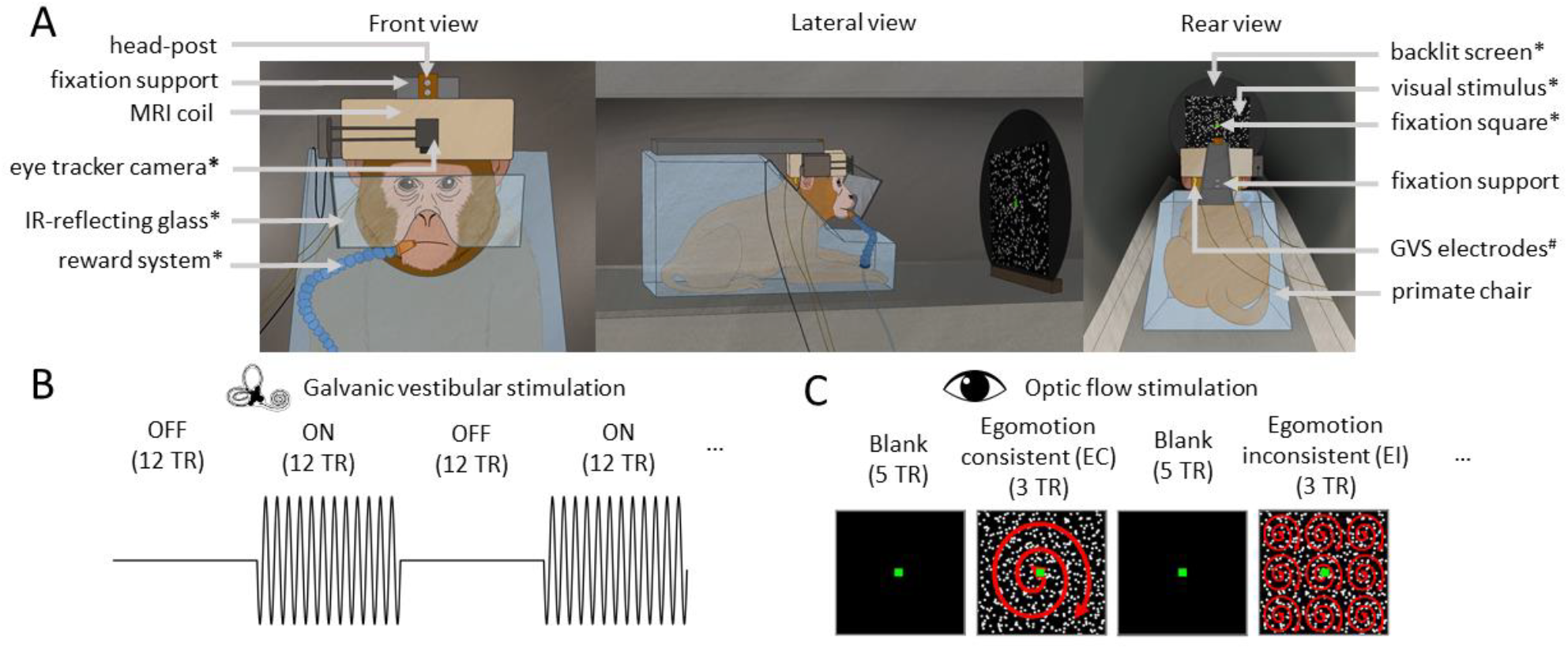
Setup and stimulation protocols for monkey fMRI experiments. (A) Schematic representation of the monkey fMRI setup. The animal is seated in a sphinx position within a primate chair, placed inside the MRI scanner bore, with the head restrained using a head-post. An 8-channel phased-array coil is positioned over the head. During the visual stimulation protocol, the monkey performs a passive fixation task by maintaining gaze on a green fixation square, back-projected onto a screen via a video projector. Eye position is monitored using an infrared video-based eye tracker. Correct fixation triggers fluid rewards throughout the runs. Components specific to the optic flow stimulation protocol are marked with an asterisk (*). In the GVS protocol, the monkey is slightly anesthetized but remains in the same head-fixation setup. Low-intensity electrical currents are delivered through gold electrodes placed behind each mastoid. Elements specific to the GVS protocol are marked with a hash (#). (B) GVS protocol. During GVS runs, animals received alternating blocks of electrical stimulation (ON) and rest (OFF), each lasting 12 TRs. Each animal completed 12 runs, with 145 volumes acquired per run. The stimulation was delivered while the animal remained head-fixed in the MRI setup under light anesthesia. (C) Illustration of the visual stimuli and block design used during optic flow stimulation. Egomotion-consistent (EC) stimuli consisted of a field of moving dots forming time-varying patterns of expansion, contraction, and rotation, consistent with self-motion along a spiral trajectory. Egomotion-inconsistent (EI) stimuli consisted of a 3 × 3 array of identical smaller EC stimuli, disrupting the global motion consistency. Runs followed a block design alternating EC and EI stimuli, interleaved with blank fixation periods. Each run comprised 7 blocks (total duration: 112 TRs). Half of the runs began with an EC block, and the other half with an EI block.

#### 2.3.2. Optic flow stimulation

The optic flow stimuli were similar to those used in earlier studies in humans (Cardin & Smith, 2010; Wall & Smith, 2008) and followed an identical protocol as the one described in Cottereau *et al*. (2017). Each stimulus was a 6-seconds video of 800 white dots moving on a black background, shown at 60 frames per second (fps). In the egomotion-consistent (EC) condition, dots moved according to a coherent flow field with expansion, contraction, and rotation, simulating self-motion along a spiral path. The egomotion-inconsistent (EI) condition consisted of a 3 × 3 grid of EC panels that disrupted the global optic flow needed to detect self-motion. Dot size, velocity, and number of dots were equalized across conditions to control for low-level visual information differences. For a full description of the motion equations, see Cottereau *et al*. (2017). The macaques were placed in the primate chair, in the sphinx position, with their heads fixed to the chair using a plastic implant attached to the surface of their skull, and with a tube dispensing water placed at the mouth entrance, connected to an automatic system, rewarding the monkey when it looked into a fixation window of about 2 × 2 degrees around the fixation target at the center of the screen. Monkeys in the primate chair were introduced into the bore of the magnet, facing a translucent screen at a distance of 50 cm. Visual stimuli were rear-projected on the screen by a video projector (Hitachi, CP_X809), at a spatial resolution of 1024 × 768 pixels and a refresh rate of 60 Hz. The position of one eye was monitored with an infrared video-based eye-tracker at 60 Hz (ASL). Throughout the acquisition, a green square (pixel size: 20 × 20 pixels, DVA size: 0.44 × 0.44, relative size: 0.02 × 0.03) was also projected in the center of the screen to maintain the monkey’s gaze. The fixation area for rewarding the monkey was a circle (pixel size: 180 × 180 pixels, DVA size: 3.93 × 3.93, relative size: 0.18 × 0.23, visible only to the experimenters from the MRI control room).

#### 2.3.3. Data acquisition

##### Anatomical and functional brain templates

The MRI images were acquired by a clinical 3T scanner (Philips Achieva), equipped with a custom 8-channel phased array coil (RapidBiomed) designed to fit the skull of our monkeys. For each individual, anatomical and functional brain templates were built from acquisitions made in a single session on slightly anaesthetized animals (Zoletil 100: 10 mg/kg and Domitor: 0.04 mg/kg), during which we acquired four T1-weighted anatomical volumes (TR = 10.3 ms, echo time [TE] = 4.6 ms, flip angle = 8°, voxel size = 0.5 × 0.5 × 0.5 mm, 192 slices), and 300 functional volumes (GE-EPI, TR = 2000 ms, TE = 30 ms, flip angle = 90°, SENSE factor = 1.6, voxel size = 1.25 × 1.25 × 1.5 mm, 32 slices).

##### GVS

Each animal participated in one session under slight anesthesia (Medetomidine: 0.04 mg/kg and Ketamine: 10 mg/kg). Each session lasted about 45 minutes, during which 12 runs were recorded. Each run lasted 216 s (145 TRs) and consisted of alternating periods of galvanic stimulation (ON) and rest (OFF). There were two types of runs, with the same temporal structure but starting either with ON or OFF conditions (types 1 and 2, respectively). During previous sessions, four T1-weighted anatomical volumes by a magnetization-prepared rapid-acquisition gradient-echo sequence (MPRAGE repetition time (TR) = 10.3 ms, echo time (TE) = 4.6 ms, flip angle = 8°, voxel size = 0.5 × 0.5 × 0.5 mm, 192 axial slices with no inter-slice gap, field of view (FOV) = 160 × 160 mm) were obtained. During the sessions, 145 T2-weighted functional volumes were acquired by gradient echo-planar imaging (GE-EPI: TR = 1500 ms, TE = 30 ms, flip angle = 75°, SENSE factor = 1.6, voxel size = 1.25 × 1.25 × 1.5 mm, 46 axial slices with no inter-slice gap, FOV = 80 × 80 mm). These T1 and EPI volumes were used to construct individual anatomical and functional templates, respectively.

##### Optic flow

Functional images of the optic flow protocol were made on behaving awake macaques. Sessions lasted a maximum of one hour per day, and were repeated on successive days until 36 valid runs were obtained, *i*.*e*. runs in which the monkey looked at least 85% of the acquisition time in the reward window of about 2 × 2 degrees centered on the fixation target. Images were acquired with the same GE-EPI sequence used during the anesthetized sessions. EC and EI stimuli were presented using a block-design built with EventIDE software (Okazolab). Each run consisted of 224 s (112 TRs) divided into 7 similar cycles of 32 s (16 TRs). In half of the runs, a cycle started with a baseline of 10 s (5 TRs) where only the fixation point was present. It was followed by 6 s (3 TRs) of the EC condition, then by another 10 s of blank and finally by 6 s of the EI condition. In the other half of the runs, the EC and EI conditions were reversed within a cycle (*i*.*e*., a cycle had 10 s of blank, 6 s of the EI condition, 10 s of blank, and finally 6 s of the EC condition) (**Figure 1C**).

### 2.4. Data Processing

#### 2.4.1. Anatomical and functional templates

To build the anatomical template, four T1-weighted (MPRAGE) images were originally realigned and averaged. The resulting volume was then registered onto the NMT v2.0 template space (Jung *et al*., 2021; Seidlitz *et al*., 2018). Functional templates were established through the averaging of 300 GE-EPI volumes after motion correction. After brain extraction of the anatomical and functional volumes, we normalized the functional volumes to the anatomical reference using SPM12 (http://fil.ion.ucl.ac.uk/spm/) non-linear normalization tools, and the deformation parameters were saved for application to the functional volumes acquired during GVS and optic flow stimulations. Another non-linear normalisation was used to register the NMT v2.0 volume to each of the individual structural templates. These normalisation parameters were used to warp the NMT v2.0 cortical surfaces and CHARM atlas into the individual space of each animal.

#### 2.4.3. Preprocessing of the functional data

*GVS*. Functional images were first converted from DICOM to NIFTI format using dcm2niigui (MRIcron) (Li *et al*., 2016). For each run, functional volumes were first motion-corrected using rigid realignment. Slice timing correction was then applied to account for acquisition delays across slices. A mean image was computed from each run and used to co-register the data to the individual’s functional template. This run-specific registration was combined with precomputed, subject-specific normalization parameters that aligned the functional template to the anatomical volume. Both sets of transformations were applied simultaneously during a single resampling step, producing volumes with 1 × 1 × 1 mm resolution and applying slight spatial smoothing using a Gaussian kernel (FWHM = 2 mm isotropic). Additionally, we applied band-pass filtering (0.01-0.25Hz) on the GVS functional time series to alleviate the artifacts potentially caused by GVS. All these preprocessing steps were performed with SPM12.

##### Optic flow

Functional images acquired during optic flow stimulation were preprocessed in the same way as those acquired during GVS stimulation except that we did not apply band-pass filtering of the functional time series.

### 2.5. Statistics and Results Presentation

#### GVS

Voxel-wise statistics were computed for each individual by fitting a general linear model (GLM) to the blood oxygen level-dependent (BOLD) signal. The model contained two main regressors representing the two experimental conditions: ON and OFF. These regressors were convolved with the hemodynamic response function (HRF) estimated from three of the four monkeys (see Cottereau *et al*., 2017). In addition, we included 6 noise regressors per run to capture acquisition and physiological noise. They were computed by submitting the time series of all the non-brain tissues (background, eye balls, masticatory muscles, etc.) to a principal component analysis (PCA) from which we retained the 6 regressors capturing the most variance (Vanduffel, 2014). Both the preprocessing steps and GLM analyses were implemented in MATLAB, with the SPM12 software and custom scripts.

#### Optic flow

The GLM included three major regressors of the experimental conditions—egomotion consistent and inconsistent conditions (EC and EI), and blank periods—each convolved with individually-estimated HRF. Noise regressors were computed with the PCA method described above. However, since the animals were awake, the noise sources were potentially more numerous and we thus included 12 noise regressors rather than 6 in the model.

For both GVS and optic flow, we performed statistical inference through cluster-extent based thresholding to correct for multiple comparisons. A voxel-wise threshold of p < 0.001 (uncorrected) was used to identify contiguous clusters and the resulting clusters were then tested against a clust er- extent threshold controlling for cluster family-wise error (FWEc) at p < 0.05 (k = 32) (Phillips, 2018). The thresholded statistical parametric maps (SPM) were then projected on the individual cortical surfaces using Connectome Workbench (https://www.humanconnectome.org/software/connectome-workbench), with the “enclosing voxel” algorithm. Since all these cortical surfaces were derived from the NMTv2.0 symmetrical cortical surfaces, surface-based results from all four monkeys (eight hemispheres) were then projected on the right cortical surface of the NMT template. Results were considered significant at the group level for all surface nodes in which the individual cluster-extent based threshold was reached in at least three out of the eight hemispheres (*i*.*e*. at least two out of the four individuals).

### 2.5. Regions of interest (ROIs) analyses

We used the surface-based CHARM atlas to perform ROIs analyses. First, for each cortical region of the CHARM atlas (n=135), we computed the percentage of surface nodes showing significant group activation (>3 hemispheres) for GVS, for optic flow, and for both sensory modalities. We retained only the regions in which at least 10% of the nodes (10% coverage) were activated for at least one sensory modality. Then, for each CHARM’s region, we computed the mean percent signal change (PSC) across all nodes for the contrasts ON >OFF (GVS) and EC>EI (optic flow). These PSC were submitted to one-tailed t-tests for each of the eight hemispheres and the resulting individual p-values were combined across the eight hemispheres using Fisher’s method. Since these statistical operations were repeated for each CHARM region, the resulting group-based p-values were then corrected for multiple comparisons using Bonferroni correction.

## 3. Results

### Cortical Network Involved in Processing Galvanic Vestibular Stimulation

In the present study, we use functional MRI to investigate the cortical areas involved in GVS processing in non-human primates. To this end, four rhesus macaques were exposed to GVS, with stimulation (ON periods) interspersed with pauses without stimulation (OFF periods) (see Materials and methods). We then observed changes in the BOLD signal between the ON and OFF conditions. **Figure 2** shows the statistical parametric maps (t-score) based on all 12 runs per animal. They are projected on inflated representations of their left and right cortical surfaces with cluster-extent based thresholding for multiple comparisons (p < 0.05 FWEc-corrected). Within these four individuals, robust activations were observed, notably in the superior temporal cortex, posterior parietal cortex, insular and opercular-insular regions, cingulate and medial cortical regions, and in at least three of the individuals, in the frontal cortex. We observe variable interhemispheric asymmetries on the activation maps of some of our individuals (*e*.*g*. M01 and M04), without showing consistent lateralization across our sample.

**Figure 2.**
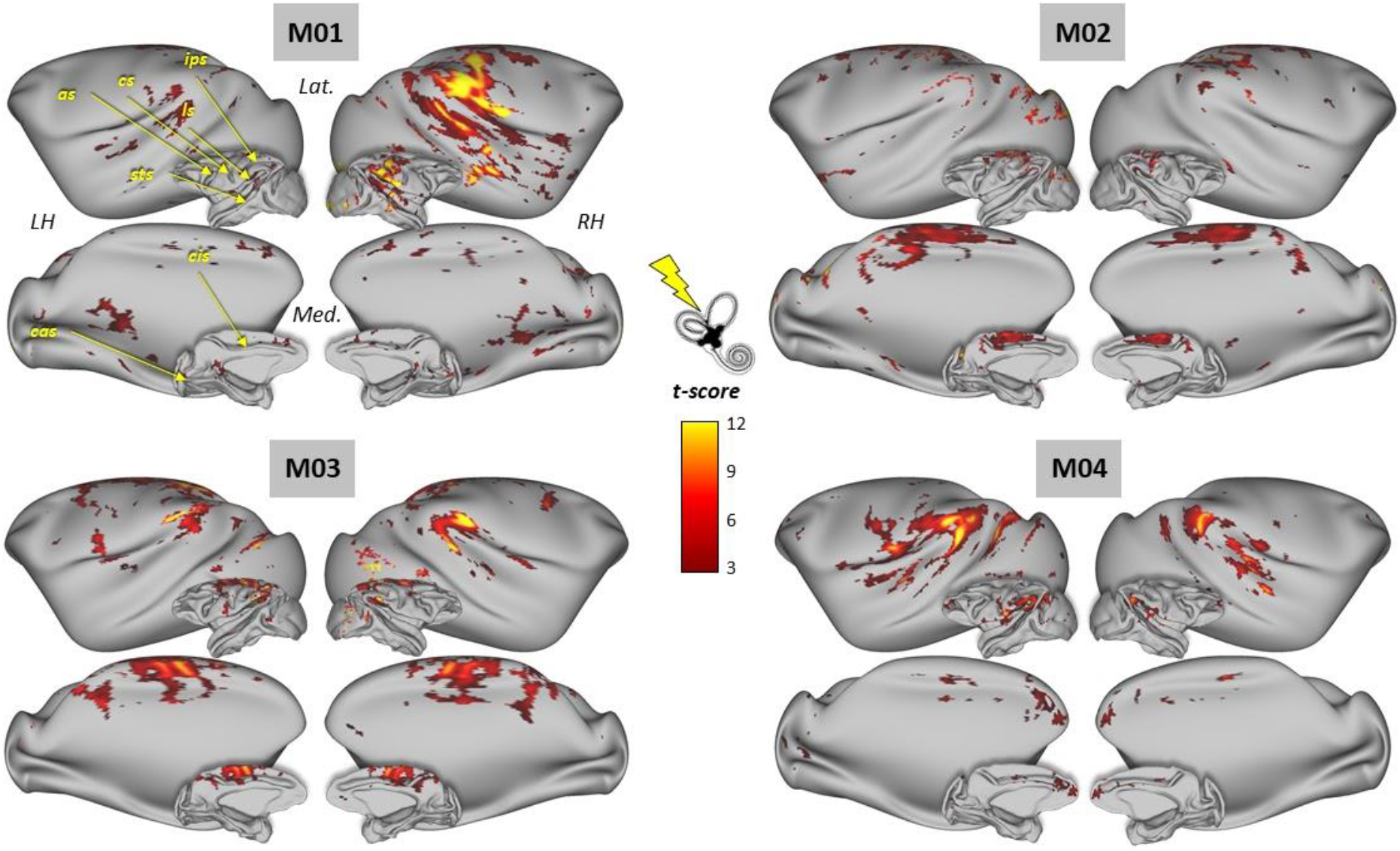
Cortical activations evoked by GVS in the four macaque monkeys. For each monkey (M01, M02, M03, M04), statistically significant BOLD activations in the contrast “GVS ON” > “GVS OFF” (cluster-extent threshold, p < 0.05 FWE-corrected) are shown on lateral (Lat.) and medial (Med.) views of both inflated and fiducial representations of the monkeys’ left and right hemispheres (LH and RH, respectively). The principal sulci are indicated in yellow in the upper left M01 cortical surfaces (*as*: arcuate sulcus; *cs*: central sulcus; *cas*: calcarine sulcus; *cis*: cingulate sulcus; *ips*: intraparietal sulcus; *ls*: lateral sulcus; *sts*: superior temporal sulcus).

### Cortical Network Involved in Egomotion-coherent Optic Flow Processing

We also used fMRI to study the cortical areas involved in egomotion-coherent optic flow processing in the non-human primate. To this end, the same four rhesus macaques were exposed to optic flow patterns that were either consistent (EC condition) or inconsistent (EI condition) with egomotion (see Materials and methods). We then observed changes in the BOLD signal between the EC and EI conditions. **Figure 3** shows parametric statistical maps (t-values) based on all available data (36 runs/animal). Optic flow coherent with egomotion was found to elicit statistically robust increases in BOLD signal, with activation clusters having t-scores ranging from 3 to 8 (cluster range threshold, FWE-corrected p < 0.05). In all four individuals, robust activations were observed, notably in the superior temporal cortex, posterior parietal cortex, medial superior temporal areas and frontal cortex, in agreement with previous studies (Cottereau *et al*., 2017; De Castro *et al*., 2021).

**Figure 3.**
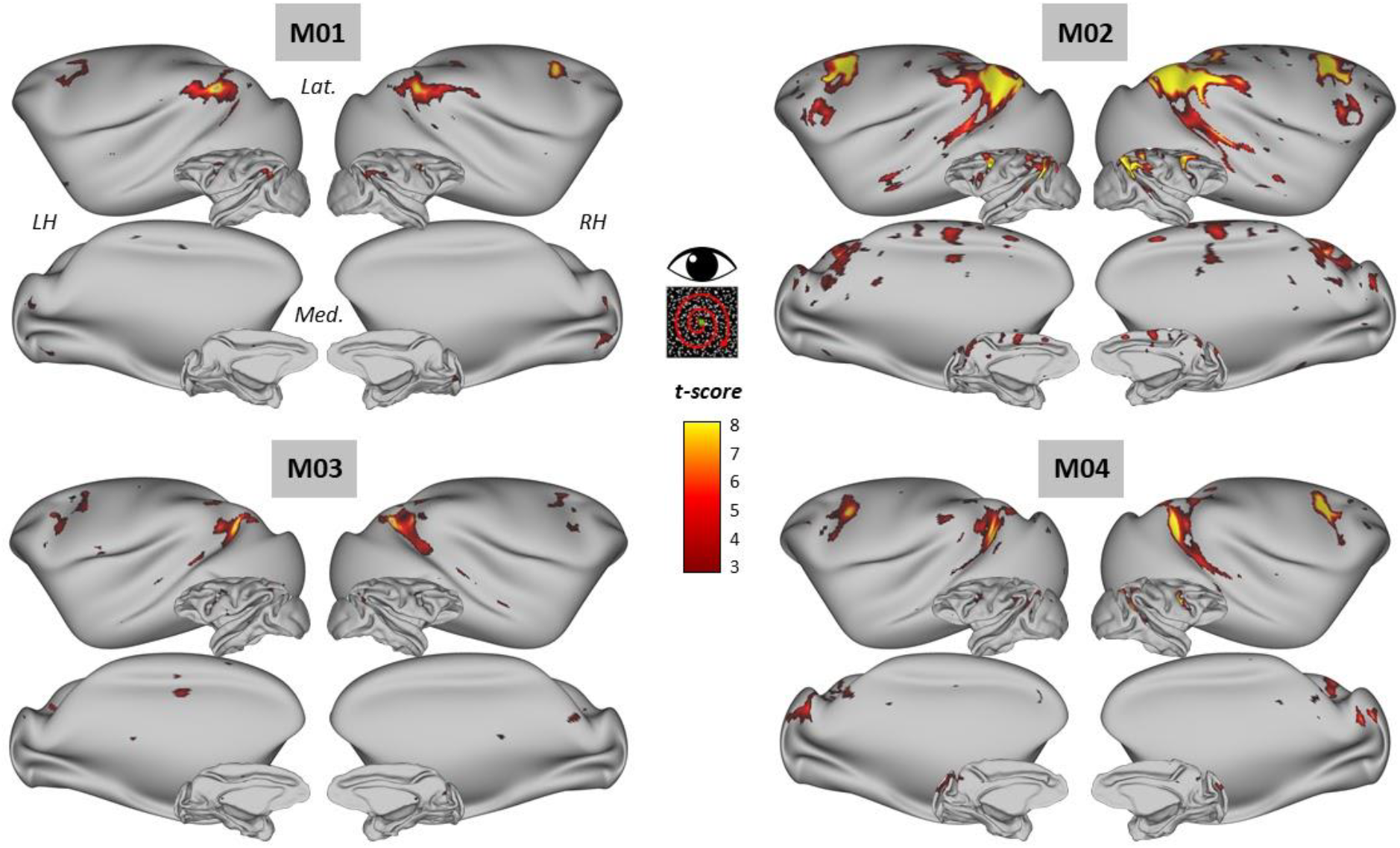
Cortical activations evoked by egomotion-consistent optic flow in the four macaque monkeys. For each monkey (M01, M02, M03, M04), statistically significant BOLD activations in the contrast “1P” > “9P” (cluster-extent threshold, p < 0.05 FWE-corrected) are shown on lateral (Lat.) and medial (Med.) views of both inflated and fiducial representations of the monkeys’ left and right hemispheres (LH and RH, respectively).

### Overlap of vestibular and visual activations

The thresholded activation maps observed for our four animals (*i*.*e*. eight hemispheres) in response to GVS were then binarised (1: significant activation, 0: no significant activation) and summed up after node-to-node projection onto the right cortical surface of the NMT v2.0 template (**Figure 4A**). This led to a new “group” map in which each surface’s node has a score between 0 and 8, depending on the number of hemispheres in which that particular node exhibited statistically significant activation. Thresholding this group map at three hemispheres (*i*.*e*. at least two significant individuals) provides a clean, extensive and robust vestibular cortical network. The most consistent regions found in at least five of the eight hemispheres include the ventral posterior suprasylvian (VPS), the parieto-insular vestibular cortex (PIVC), the primary and secondary somatosensory cortices (S1/2) and the primary motor cortex (M1). Extending to regions present in at least three of the eight hemispheres, *i*.*e*. present in at least two of our four individuals, we also find the dorsal subdivision of the medial superior temporal area (MSTd), the parietal area 7 (7), the ventral intraparietal area (VIP), the granular insular area (Ig), the belt portion of the auditory cortex (BELT), the superior parietal lobule (PE), the dorsal caudal premotor cortex (PMdc), the smooth eye movement subdivision of the frontal eye field (FEFsem), the cingulate sulcus visual area (CSv), the pre-supplementary motor area (PreSMA) and a portion of the precuneus gyrus (PGm).

**Figure 4.**
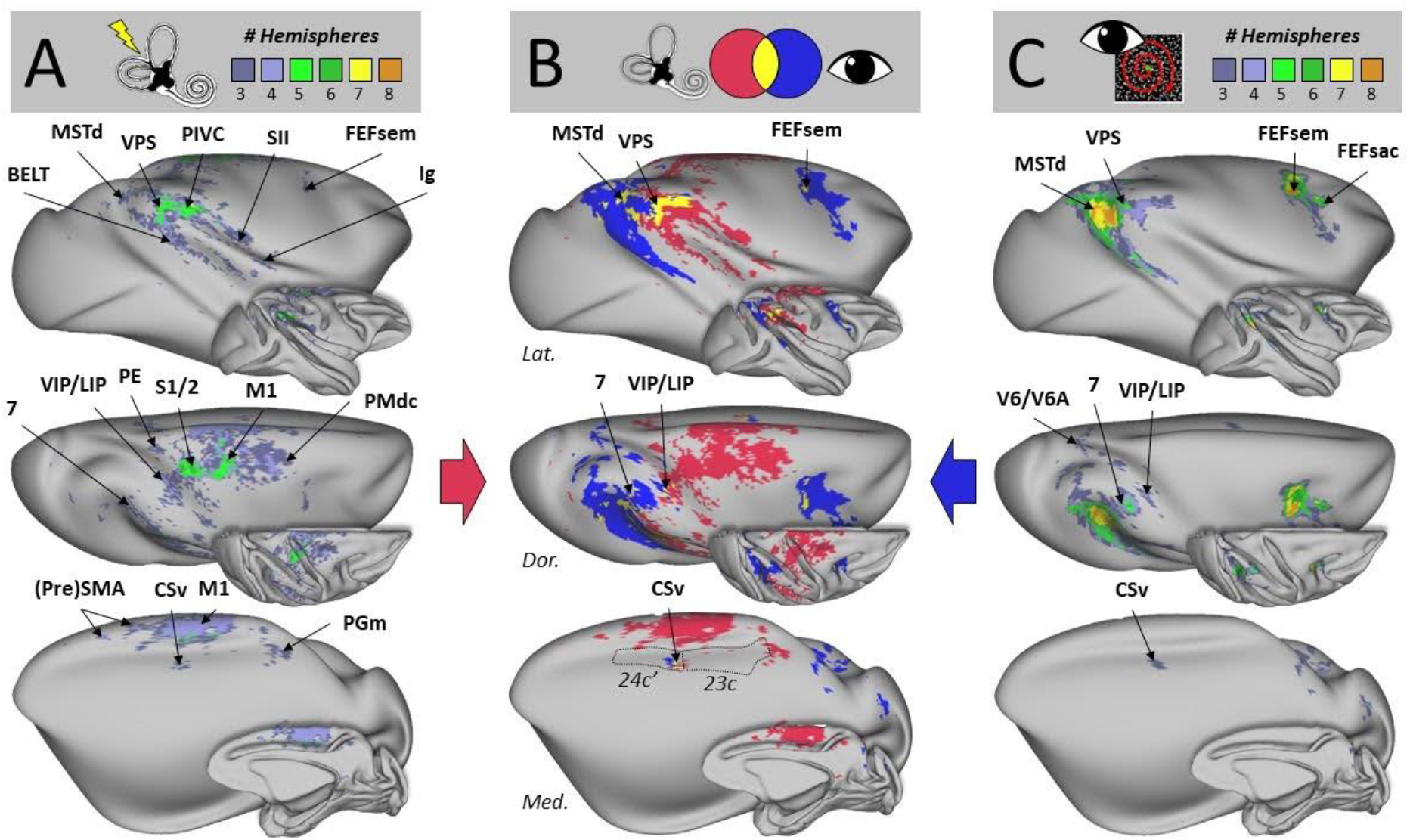
Group-level cortical responses to vestibular and visual egomotion stimuli. (A) Cortical activations evoked by GVS at the group level. Results for the left and right hemispheres of the four individual monkeys (*cf*. Figure 2) are superimposed on lateral (Lat.), dorsal (Dor.) and medial (Med.) views of both inflated and fiducial representations of the right cortical surface of the NMT template. Only activations found in at least three hemispheres (*i*.*e*. at least two individuals) are shown. (B) Overlap of vestibular and visual activations. Activations evoked at the group level by GVS or egomotion- consistent optic flow are shown in magenta and blue respectively, while the regions responding to both types of stimuli are indicated in yellow. Dotted lines in the medial view (lower panel) illustrate the borders of the cingulate areas 24c’ and 23 in which CSv lies. (C) Cortical activations evoked by egomotion consistent optic flow stimulation at the group level. Same conventions as **Figure 4A**.

By proceeding identically with acquisitions in the visual modality, **Figure 4C** reveals a well-known network for the processing of visual self-motion information (*i*.*e*., a constant presence in each of the eight hemispheres of MSTd and FEFsem), presence in the majority of hemispheres (at least four out of eight) of parietal area 7, and presence in at least two of the four individuals (at least three out of eight hemispheres), of VPS, the saccadic subdivision of the frontal eye field (FEFsac), V6/V6A, VIP and CSv.

Consequently, an extensive and robust network involved in processing egomotion related information is observed for each of these sensory modalities. However, as shown in **Figure 4B**, the vestibular network (in magenta) and the visual one (in blue) are predominantly distinct from each other. The vestibular network extends from the temporo-parietal junction (tpj) to the tip of the superior temporal sulcus (sts) along the length of the lateral sulcus, coupled with medial activations in the motor and premotor regions. By contrast, the visual network occupies a territory from the tpj to the more posterior portions up to the occipito-parietal junction, coupled with a large frontal patch located at the level of the frontal eye field (FEF). Despite the relatively large extent of the vestibular and visual cortical networks, the sites of visuo-vestibular convergence (in yellow in **Figure 4B**) are only found in a restricted number of areas, namely MSTd, VPS, area 7, FEFsem, VIP, LIP and CSv.

### Quantification of the visuo-vestibular convergence

In order to situate the results shown in **Figure 4B** with respect to a well-established atlas space, we used the CHARM atlas (Reveley *et al*., 2017; Jung *et al*., 2021) to calculate, for each of the 135 atlas regions, the percentage of cortical nodes (% coverage) showing significance for the vestibular modality only, for the visual modality only and for both modalities. In order to focus on regions with substantial cortical involvement while excluding minor, potentially spurious activations, we only considered atlas regions presenting at least 10% of activated nodes for at least one modality. Results of this analysis are shown in Figure 5A as bar plots of percent coverage for the thirty-three out of 135 atlas regions meeting the above-mentioned criteria. In response to GVS, twenty regions show significant coverage (magenta bars), including regions of the parietal cortex (LIPv, LIPd, VIP, 7op, 7b, 7a, PE, PEa), premotor cortex (PMdc, SMA), sensorimotor cortex (S1/2, S3a/b, M1), temporal cortex (MST, Tpt), as well as the caudo-lateral, medio-lateral and caudo-medial portions of the auditory belt cortex (BCL, BML, BCM respectively, and insular areas (AI, Ig). Using the same approach for EC optic flow stimulation, twenty-two regions showing significant coverage (blue bars) are identified, covering the visual cortex (MT, MST, FST, V4d, V3a, V6Ad), parietal cortex (LIPv, LIPd, VIP, MIP, LOP, 7op, 7b, 7a, PGa), premotor and motor cortex (PMdc, F5), frontal cortex (44, 45b, 8Bs, 8Ad), and the temporoparietal junction (Tpt). Confirming the visual impression given by Figure 4B regarding the moderate overlap between the vestibular and visual networks, only nine out of the thirty-three regions mentioned above exhibited both vestibular and visual activations (Tpt, MST, 7op, 7b, 7a, LIPv, VIP, PMdc, LIPd). Moreover, for three of these nine regions (VIP, PMdc, LIPd), we did not observe visuo-vestibular convergence at the node level, and such convergence, in terms of nodes’ coverage, is often moderate among the six other visuo-vestibular regions (Tpt: 21%; MSTd: 16%; 7op: 14%; 7a: 10%; 7b: 7% and LIPv: 3%). Thus, this atlas-based analysis confirms that visuo-vestibular convergence is located in very specific parietal and temporal regions, with in particular the most quantitatively extended overlap found at the temporo-parietal junction (MST, Tpt, 7a), followed by the inferior parietal lobule (7b, 7op), intraparietal sulcus (LIPv, VIP, LIPd), and finally the dorsal premotor cortex (PMdc).

**Figure 5.**
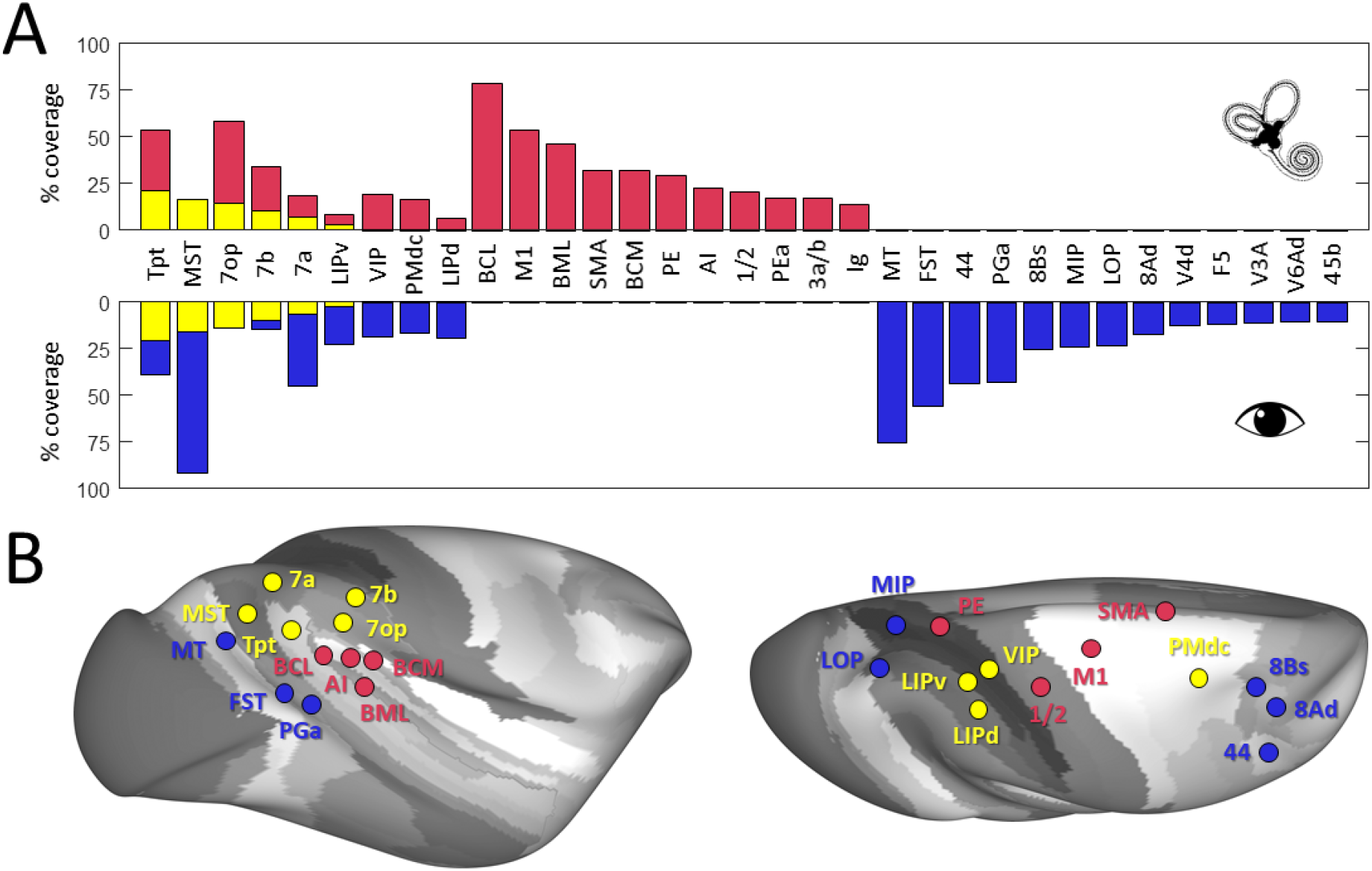
ROI analyses of the vestibular and visual convergence with respect to the CHARM atlas. (A) For each area of the CHARM atlas (n=135), we calculate the percentage of nodes exhibiting significant activations (% coverage) for vestibular stimulation (in magenta), visual stimulation (in blue) or both (in yellow). Only areas showing at least 10% of activated nodes for one sensory modality are shown as bar plots. (B) The location of these CHARM areas is shown in lateral and medial views of the inflated right hemisphere of the NMT template. Vestibular and visual areas are shown in magenta and blue respectively, while areas exhibiting activations for both modalities are reported in yellow.

By superimposing the vestibular and visual networks mapped in the same four monkeys (**Figure 4B**), we could precisely identify seven visuo-vestibular areas: MSTd, area 7, VPS, FEFsem, VIP, LIP and CSv. The atlas-based analysis (**Figure 5**) largely confirms and extends these findings. Tpt (that includes VPS) and MST (that includes MSTd) are the two CHARM areas showing the greatest visuo-vestibular overlap. They are followed by parietal area 7 which, in the CHARM atlas, comprises three regions: 7a, 7b and 7op (that latter includes PIVC as it is the case for its human homologue; Grüsser *et al*., 1990; Lopez *et al*., 2012). Interestingly, these three visuo-vestibular regions exhibit a gradient in which 7a, posteriorly, is dominated by visual activations and 7op, anteriorly, is dominated by vestibular activations, while 7b, in between, responds about equally well to both sensory modalities. The atlas based-analysis also shows that both the ventral and dorsal subdivisions of LIP (LIPv and LIPd respectively) are activated by visual and vestibular inputs. However, as it is also the case for VIP and PMdc (that includes FEFsem), the within-area node-to-node correspondence between the vestibular and visual activations is weak or absent. The only missing area in the atlas-based analysis is the most recently discovered area CSv, that lies within the cingulate sulcus. CSv is not reported in the CHARM atlas, where it is diluted between the large cingulate areas 24c’ and 23c, as illustrated in the lower panel of **Figure 4B**. Area 24c’ contains most of CSv in its posterior portion, but both the vestibular and visual coverages in this CHARM area do not reach our 10% coverage threshold (GVS coverage: 6.3%, FLOW coverage: 6.8%, GVS & FLOW coverage: 1.8%).

As a complement, we also used the CHARM atlas to perform a more conventional ROI analysis, computing the mean BOLD PSC for GVS and optic flow for each of the atlas regions and in each individual. p-values obtained in the four individuals (eight hemispheres) were combined using Fisher’s method and corrected for the number of CHARM regions (Bonferroni correction, pFWE < 0.05). The results of this ROI analysis are shown in Figure 6. Twenty-four areas showed statistically significant responses to GVS alone (red symbols), nine areas responded significantly to optic flow alone (blue symbols), and importantly, ten areas demonstrated significant bimodal responses (yellow symbols, p <

**Figure 6.**
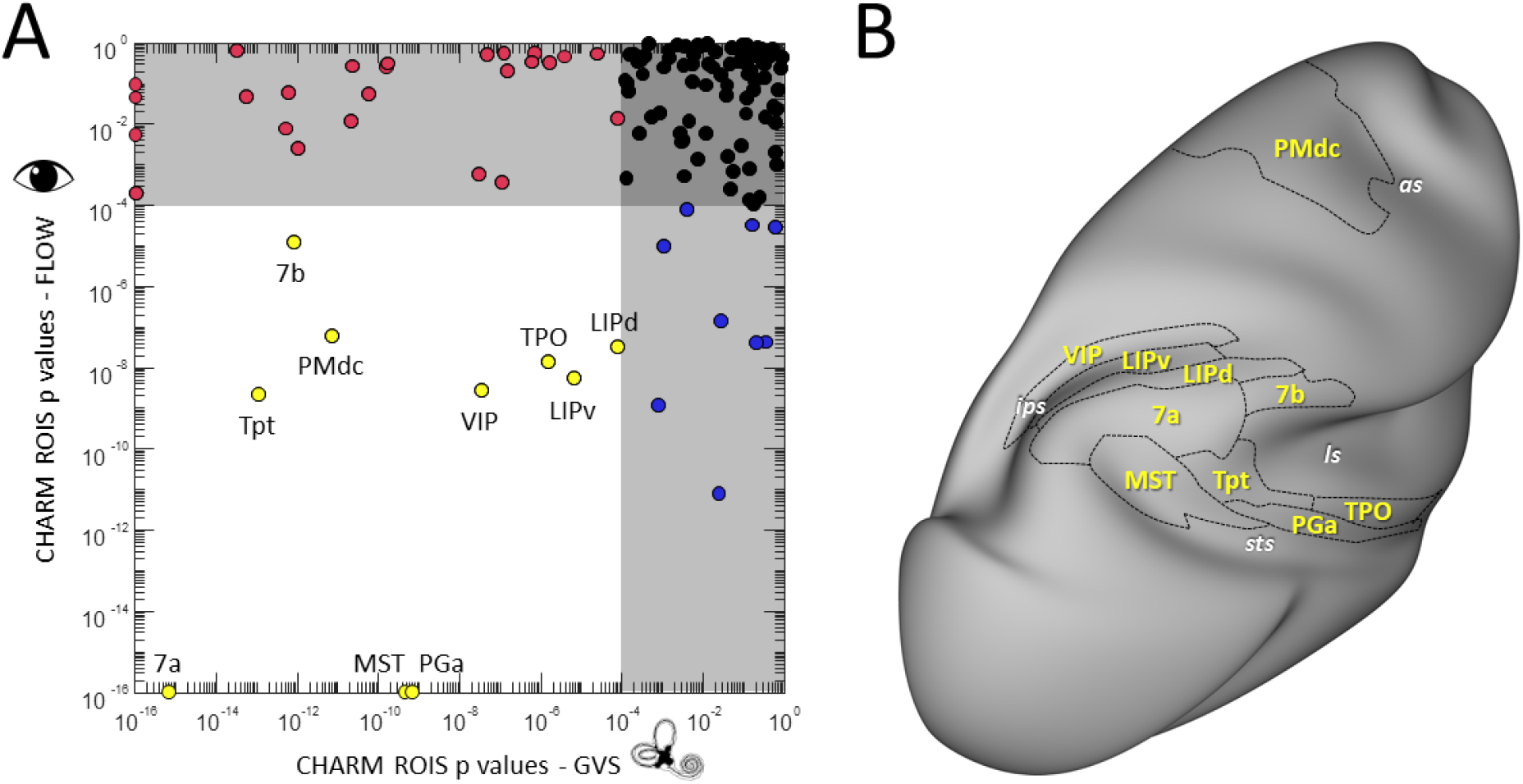
ROI analyses of the vestibular and visual activations with respect to the CHARM atlas. (A) For each area of the CHARM atlas (n=135) and each monkey, we calculated the p-value for the vestibular (“GVS ON” > “GVS OFF”) and visual (“1P” > “9P”) contrasts (see Methods section). For each modality, individual p-values were combined using Fisher’s method. Combined p-values for GVS and FLOW are shown for the 135 CHARM areas (circular symbols). Black, magenta, blue and yellow symbols stand for areas statistically non-significant for both GVS and FLOW, significant for GVS only, significant for FLOW only and significant for both GVS and FLOW, respectively (p < 0.05 after Bonferroni correction for the number of CHARM areas). (B) The borders of the CHARM areas significant to both GVS and FLOW are shown on an inflated representation of the right hemisphere of the NMT template.

0.05 after Bonferroni correction). These regions comprise the areas previously mentioned in the other analyses: parietal area 7 (7a/7b), Tpt (VPS), PMdc (FEFsem), MST (MSTd), LIP (LIPd/LIPv) and VIP. In addition, this statistically more powerful approach reveals two additional areas, the temporo-paerieto- occipital area (TPO) and PGa, both lying in the upper bank of the sts, and thought to be parts of the superior temporal polysensory (STP) area (Bruce *et al*., 1981; Seltzer & Pandya, 1994).

## Discussion

The present study aimed at characterizing the cortical networks involved in processing vestibular and visual signals generated by egomotion in non-human primates. Using fMRI in four macaque monkeys, we investigated cortical responses to GVS and optic flow patterns consistent with egomotion. Our results indicate that egomotion-related vestibular and visual stimuli are handled by widely dissociated cortical networks. GVS activates preferentially vestibular, associative and, motor/premotor areas located in the insular, superior parietal, temporal, frontal and cingulate cortices. We also observed robust vestibular activations in the auditory cortex, mostly in its belt portion, which is coherent with the fact that the belt is fed with somatosensory information, notably through anatomical connections with insular areas (Kayser *et al*., 2005; Smiley & Falchier, 2009). By contrast, optic flow elicits preferentially occipito-temporal, inferior parietal and prefrontal oculomotor regions and the cortical network documented here has been described with more details in previous studies (Cottereau *et al*., 2017; De Castro *et al*., 2021). However, some areas of convergence are highlighted, where both modalities elicit strong activation. Statistical testing shows that seven regions in total show strong responses to both sensory modalities with widespread cortical coverage: MSTd in the superior temporal sulcus, VPS at the posterior tip of the lateral sulcus, CSv within the cingulate sulcus, FEFsem within the frontal eye field region, area 7 in the inferior parietal lobule, and both LIP and VIP within the intraparietal sulcus. Beyond these primary convergence sites, our analyses also reveal additional regions showing consistent responses to either modality and, in some cases, both, including subdivisions of area 7 in the inferior parietal lobule (*i*.*e*. 7op, 7b), subdivisions of the lateral intraparietal area (LIPv and LIPd), and premotor dorsal caudal cortex (PMdc). These results indicate that visuo-vestibular integration arises from selective convergence within a few specific cortical locations rather than from a general overlap of vestibular and visual networks, providing new insight into the neural mechanisms underlying multisensory processing of self-motion information in primates. **Figures 4 and 5** show a high degree of consistency, especially when it comes to overlapping regions like 7a, VIP and MSTd. Several other areas, like FEFsem and PMdc, appear to represent adjacent or functionally related territories despite having different labels. This convergence across statistical approaches supports the robustness of the identified network involved in processing vestibular and visual cues for egomotion perception.

Our results confirm that visuo-vestibular convergence occurs within a restricted set of areas. Notably, our labeling of VPS, MSTd, area 7, LIP, VIP, FEFsem and CSv as convergence regions aligns with earlier single-unit and fMRI findings that these regions are involved in integrating visual and vestibular information for egomotion (Chen *et al*., 2011b, 2011c, 2013; Gu *et al*., 2006, 2008, 2010; Schlack *et al*., 2002; Smith *et al*., 2012).

In our study, VPS (Tpt) is the area showing the greatest overlap between vestibular and visual activations, with a slight dominance of vestibular modality (*cf*. **Figure 5**). VPS is located posterior to PIVC (7op) and, like that latter, it has been shown to receive short-latency vestibular input (Akbarian *et al*., 1992). However, it also receives input from a portion of the sts likely encompassing MSTd (MST) (Guldin & Grüsser, 1998). The slight vestibular dominance documented here is consistent with previous electrophysiological studies showing that the responses of visuo-vestibular neurons in VPS are often dominated by the vestibular modality, with generally opposite directional tuning between both sensory modalities (Chen *et al*., 2011b).

The second area of greatest visuo-vestibular overlap documented here is MSTd (MST). Previous studies showed with great rigor the distinction between the dorsal (MSTd) and ventral (MSTv) parts of the MST area, evidencing that MSTd mainly processes optic flow information for the perception of self-motion, while MSTv processes more object motion (Tanaka & Saito, 1989). MSTd is therefore a well documented cortical area for egomotion processing of visual input (Duffy & Wurtz, 1991; Saito *et al*., 1986; Tanaka *et al*., 1986). This region is responsive to EC optic flow in our data set, and its GVS response was less significant, which is in line with previous findings showing that visual responses are dominant in this area (Gu *et al*., 2006; Morgan *et al*., 2008; Takahashi *et al*., 2007). While it belongs to this bimodal network, MSTd cells typically have congruent or reverse visual/vestibular tuning for translation but reverse preferences for rotation (Takahashi *et al*., 2007), which suggests specialization for solving self-translation signals (Gu *et al*., 2006). This is consistent with our results indicating that MSTd is a visuo-vestibular convergence point with visual bias (*cf*. **Figure 5**).

Our results indicate that the third region showing greatest visuo-vestibular overlap is the parietal area 7. The subdivision of area 7 is a matter of debate. However, regardless of the nomenclature system used, the properties of sensory specialization remain consistent. With regard to the initial subdivision, area 7a of the superior posterior parietal cortex has been identified as essential for visuo-vestibular integration in primates (Andersen, 1997; Snyder *et al*., 1998) and is strongly activated by optical flow in humans and macaques (Claeys *et al*., 2003; Deutschländer *et al*., 2002). Traditionally, 7b has been considered a secondary somatosensory area involved in the processing of tactile and proprioceptive information (Andersen, 1997). However, visual and bimodal (visuo-tactile, visuo-cochlear) neurons have been described (Fogassi *et al*., 2005; Hyvärinen, 1981). Some subregions of 7b are close to the vestibular areas identified in the parietal operculum, but 7b itself is not classically described as a pure vestibular area. As for 7op, it is a region close to or including areas identified in macaques and humans as being involved in vestibular processing, notably PIVC (Grüsser *et al*., 1990; Lopez *et al*., 2012). However, this traditional subdivision into portions 7a and 7b has been extended by further anatomical studies. Cytoarchitectonic and connectional data (Rozzi *et al*., 2006) suggest that area 7a actually comprises at least two distinct subregions, Opt and PG, with different connectivity profiles: Opt is more strongly linked to visual and oculomotor areas, while PG integrates both visual and somatosensory inputs. Similarly, the classic area 7b can be subdivided into PFG and PF, which have denser connections with premotor and somatosensory areas and are more involved in motor functions. There is therefore a clear rostrocaudal gradient, highlighted in the present study as well, along the inferior parietal convexity (IPL), with the most caudal portions (7a or Opt/PG) being more visual, the intermediate portions (7b or PG) being more equally driven by visual and vestibular input, and the rostral portions (7op or PFG/PF) showing more vestibular responses and associated with premotor areas.

VIP is a functionally defined area in the intraparietal sulcus (Colby *et al*., 1993; Duhamel *et al*., 1998). This area also turning out to be an overlap region equally activated by both modalities in the present study aligns with past studies that showed that VIP integrates vestibular and optic flow information with equilibrated sensitivity (Chen *et al*., 2011c; Schlack *et al*., 2002), and that it receives input from both MSTd (visual information) and vestibular-related areas such as PIVC (Lewis & Van Essen, 2000). VIP neurons also exhibit congruent and opposite tuning for visual and vestibular stimuli, and their combined responses can be dominated by either modality (Chen *et al*., 2011c), suggesting flexibility according to task demands. Moreover, vestibular response latency of VIP is quicker compared to MSTd (Chen *et al*., 2011a), and microstimulation of VIP may elicit optic flow-based heading judgment bias (Zhang & Britten, 2011), validating its causal role in self-motion perception.

The LIP region is functionally divided into LIPd on one side, which is mainly involved in oculomotor planning, and LIPv on the other side, involved in attentional as well as oculomotor processes (Liu *et al*., 2010). While LIP neurons do show robust responses to motion vision and vestibular stimulation, particularly for heading discrimination tasks (Chen *et al*., 2013; Gu *et al*., 2006), this area does not have more regular multisensory integration than specialized regions like MSTd or VIP. Findings suggest that LIP can serve as an important interface between multisensory self-motion signals and eye movement control (Andersen & Cui, 2009), though its integrative capacity for visuo-vestibular computation is limited relative to core vestibular areas.

Our finding of convergence in FEFsem, a region associated with eye movement control and processing of visual motion (Bremmer *et al*., 2002), is consistent with the hypothesis that visuomotor areas are activated in a modality-independent manner to regulate spatial orientation. FEFsem and PMdc are two separate but adjacent areas in the arcuate sulcus; FEFsem is located in the fundus of the arcuate sulcus, while PMdc is more caudal (Leichnetz, 2001; Tanaka & Lisberger, 2001). Despite their anatomical differences, both regions participate in complementary but distinct oculomotor functions, indicating a possible functional correspondence. While less highlighted in previous vestibular work, its presence as a bimodal region in this study is consistent with recent research in humans where FEF subdivisions are implicated in the integration of egomotion signals (Cottereau *et al*., 2017; Smith *et al*., 2018).

Finally, CSv, a visual motion- and motor control-related area in both humans (Smith *et al*., 2012; Wall & Smith, 2008) and monkeys (Cottereau *et al*., 2017), also appears on our overlap maps (*cf*. **Figure 4B**). The fact that CSv does not show up in the ROIs analyses is caused by the fact that in the CHARM atlas, this small cortical region belongs to the large cingulate sensorimotor areas 24c’ and, to a lesser extend, 23c, as illustrated in **Figure 4B**. While CSv has been shown to have robust responses to optic flow and artificial vestibular stimulation in humans (Billington & Smith, 2015; Pitzalis *et al*., 2020), decoding analysis reveals less integration in this area compared to hVIP or PIC. Our demonstration of macaque CSv convergence presents a new view, suggesting that it may serve as a sensorimotor interface to process full-field self-motion information (De Castro *et al*., 2021; Smith, 2021), though its integrative function remains to be fully described.

Some methodological limitations must be acknowledged. First, our GVS paradigm lacked a sham or tactile control condition, meaning that we cannot completely rule out the possibility that some of the observed cortical activations were driven by somatosensory rather than vestibular inputs, especially given that GVS necessarily engages vestibular afferents and cutaneous receptors at the sites of stimulation (Ferre *et al*., 2013; Stephan *et al*., 2005). Subsequent research will need to have an appropriately matched somatosensory control in order to unmistakably disentangle these contributions. Second, the GVS sessions were performed under slight anesthesia, in contrast to the awake condition used in the optic flow part of the experiment. Yet data still indicated robust and spatially coherent patterns of activation in animals, in keeping with known vestibular circuits. The anesthesia was necessary in the present study because of technical and ethical constraints of GVS delivery in macaques. Indeed, to ensure a safe behavioral response in macaques upon receiving stimulation, we believe that many months of training and study of reliable behavioral cues would be required to ensure that the animal receives stimulation at a suitable threshold so that it does not experience any pain or discomfort. Such a habituation protocol is not yet built, but it is already contemplated with the appropriate ethical considerations. Therefore, awake, trained monkey experiments could potentially be used in the future to validate and expand these results, possibly with improved spatial and temporal resolution. Finally, to directly link cortical activation with perceptual processing, it will be important to develop behavioral paradigms that assess the animals’ perception of vestibular stimulation—such as by tracking eye movement responses (*e*.*g*. torsional or vergence shifts) or through operant conditioning tasks. While such tools remain to be implemented, the convergence of our findings with previous electrophysiological and human fMRI studies supports the validity of the observed responses and reinforces the notion of a conserved cortical vestibular system across primates.

In summary, this research presents the first whole-brain fMRI mapping of cortical responses to both galvanic vestibular stimulation and egomotion-consistent optic flow in non-human primates. Through the localization of selective convergence zones in both modalities, we provide new insights into the multisensory organization of self-motion processing in the primate brain. These findings form the basis of follow-up experiments on visuo-vestibular integration dynamics in awake, behaving monkeys, enabling a better understanding of these networks’ perceptual and functional contributions.

## Data and Code Availability

The data and codes used for data analysis are available to interested parties on request from one of the corresponding authors.

## Author Contributions

Conceptualization: V.D.C., B.C., A.S.C., J.-B.D. Funding: A.S.C.

Project supervision: A.S.C., J.-B.D. Programming: M.R.

MRI acquisitions: V.D.C., E.E., M.-A.L., N.V., B.C., J.-B.D.

MRI data analysis: S.M., J.-B.D. Figure design: S.M., J.-B.D.

Writing (original draft): S.M., J.-B.D.

Review and editing: S.M., E.E., V.D.C., M.-A.L., A.S.C., J.-B.D.

## Acknowledgements

We would like to thank Marie Mercier, Allison Rodriguez, Céline Martinez and Marie Isnardon for animal care; and Frédéric Brouillet, Jean-Pierre Desirat and Sandrine Ferries for assistance with MRI acquisitions.

